# Variable zinc concentrations among commercially available Mueller-Hinton Broth brands affects SIR interpretations during broth microdilution *in vitro* antimicrobial susceptibility testing with the novel metallo-beta-lactamase inhibitor APC148

**DOI:** 10.64898/2026.06.04.730127

**Authors:** Veronika Smith, Ole Andreas Økstad, Pål Rongved

**Affiliations:** Section for Pharmacology and Pharmaceutical Biosciences, Department of Pharmacy, University of Oslo, Norway; Adjutec Pharma AS, Oslo, Norway

## Abstract

The well-known discrepancy between the *in vitro* and *in vivo* efficacy of β-lactam antibiotics in metallo-β-lactamase (MBL) containing Gram-negative bacteria has during recent years been found to be at least partially explained by zinc levels at infection sites being much lower than those found in conventional cation-adjusted Mueller-Hinton Broth (caMHB) media used for *in vitro* susceptibility testing. Previous studies have also demonstrated that a high variability exists with respect to zinc content in caMHB from different manufacturers, potentially leading to differences in SIR interpretations for β-lactam antibiotics with MBL-carrying isolates, depending on the brand of caMHB used for testing. APC148 is a zinc-chelating compound acting as an inhibitor of MBL enzymes and is currently undergoing phase one in clinical trials.

In this study, ten clinical isolates of *Klebsiella pneumoniae, Escherichia coli, Pseudomonas aeruginosa* and *Acinetobacter baumannii* harbouring MBLs (NDM, *n* = 6; VIM, *n* = 4; IMP, *n* = 1) were tested in a broth microdilution assay with meropenem and APC148, employing caMHB of various brands. One *K. pneumoniae* strain carrying only a serine-β-lactamase (KPC-2) was included as control.

Antimicrobial susceptibility testing (AST) by broth microdilution was performed according to the European Committee on Antimicrobial Susceptibility Testing (EUCAST). MICs of meropenem alone and in combination with inhibitors were tested in four cation-adjusted Mueller Hinton II (caMHB).The four caMHBs used were analysed by ICP-MS/MS and found to have highly vaiable zinc content (in the range 0.4 to 1.9 µg/mL, corresponding to 5.6 to 29.4 µM).

At 16µg/ml APC148 lowered the MIC in nearly all strains and in caMHBs from all manufacturers. At lower concentrations of APC148 (4 or 8 µg/mL), this MIC reduction could however only be retained when the zinc concentration in the broth was low, indicating that higher concentration of inhibitor is needed during *in vitro* Mic testing when using caMHB from certain manufacturers.

The present work clearly shows that not taking the zinc concentration of the caMHB used into consideration when estimating MIC performance of compounds functioning through interactions with zinc may be a considerable source of error, and specifically when investigating potential inhibitors of metallo-β-lactamase (MBL) enzymes. The present work supports the call for standardising zinc content in caMHB to be used for this purpose, to ensure that MIC results for drug combinations involving the use of zinc-chelating compounds are consistent and reproducible across laboratories.

## Introduction

In recent years, several studies have put focus on cases of observed discrepancy between the predicted efficacy of *β*-lactam antibiotics toward metallo-β-lactamase (MBL) carrying isolates by *in vitro* susceptibility testing and clinical outcome (*in vivo*). Numerous clinical reports have shown that despite results from *in vitro* MIC determination indicating resistance to carbapenem antibiotics, the same isolates were eliminated during treatment using the same drugs in patients (1–6). Similar results have been observed for other *β*-lactam antibiotics, e.g. such as cephalosporins (3, 7, 8). This effect was previously thought to be due to fitness cost, differences in infection types and patient immune responses, or misdiagnosis (9, 10). The activity of MBL enzymes is dependent on one or two zinc ions located in the enzyme active site. Under zinc deprivation, the MBL enzyme not only becomes inactive, but new enzymes synthesized under these conditions are degraded in the periplasm, possibly due to improper protein folding (11). There are also a range of examples of well-established compound classes shown to have interactions with zinc in key enzymes like the PBPs in the bacterial cell wall-building machinery, for instance the cyclic boronate class (CBOs) (12).

The majority of the zinc in the human body (99.9%) is found within human cells and cannot readily be accessed by bacterial pathogens (13). Additionally, the zinc available in plasma is mainly protein-bound, and during infection the immune system contributes to sequestering trace minerals by the host organism, thereby reducing the zinc concentration even further (13, 14). *In vivo* studies in mice have shown that at the infection site zinc concentrations are below the limit of detection by ICP-MS analysis (15). This is in stark contrast to zinc levels in commercial cation-adjusted Mueller Hinton II broth (caMHB), the standard growth medium used in *in vitro* susceptibility testing, which are not only generally considerably higher than plasma zinc levels, but also can vary considerably between manufacturers (16, 17).

APC148 is a zinc-chelating compound currently undergoing clinical trials, and which was previously shown capable of re-sensitising >98% of a collection of 234 carbapenemase-resistant MBL-positive clinical isolates of *K, pneumoniae* and *E. coli* (18). APC301 denotes the triple combination of meropenem, avibactam and APC148. A recent study employing APC301 with avibactam fixed at 8 µg/ml and APC148 fixed at 16 µg/ml showed that out of 176 MBL and SBL containing *Enterobacterales* isolates tested, all were rendered susceptible to meropenem (19).

In the present work we have performed broth microdilution MIC experiments with meropenem in combination with APC148, using cation-adjusted Mueller-Hinton Broth (caMHB) from four different manufacturers, and concomitantly assayed each batch of caMHB medium for the concentration of zinc. MIC values for the same bacterial isolate exhibited considerable variation in the different batches of media from the four manufacturers, and were dependent on the zinc concentration identified in each medium batch – and MIC variability was found to affect SIR interpretations (susceptible (S), susceptible, increased exposure (I) or resistant (R)) in accordance with the EU-CAST clinical breakpoint table (20) for the drug-BLI combination, depending on the brand of caMHB used for testing.

## Materials and methods

### Bacterial isolates

Ten MBL-producing clinical isolates (NDM, *n* = 5; VIM, *n* = 4; IMP, *n* = 1) comprising *K. pneumoniae* (n=4), *E. coli* (n=4) and *P. aeruginosa* (n=2) were included in this study. Additionally, one isolate (*K. pneumoniae* BAA1705) producing serine-β-lactamase KPC-2 as its sole β-lactamase was included as a negative control.

### Measurement of zinc concentration in Mueller-Hinton growth medium from different manufacturers

Zinc concentrations were measured in triplicate for each broth from the four manufacturers of caMHB. All samples were analysed by ICP-MS/MS using an Agilent 8900 ICP-MS/MS equipped with a multi-quadrupole ICP-MS system with collision/reaction cell technology, by SINTEF, Norway.

### Antimicrobial susceptibility testing (AST)

Antimicrobial susceptibility testing (AST) by broth microdilution was performed according to the European Committee on Antimicrobial Susceptibility Testing (EU-CAST) Reading guide for broth microdilution (Version 5.0, 2024; http://www.eucast.org). Minimum inhibitory concentrations (MICs) for meropenem alone and in combination with inhibitors were performed in triplicate, and modal MICs were reported. AST testing was performed for all strains (Table 2) in cation-adjusted Mueller Hinton II (caMHB) broths from four different manufacturers (Oxoid, CM0405B; Becton Dickinson (BD), 212322; Millipore, 90922; Thermo Scientific (Thermo Fischer), T3462). The assays were performed in premade Sensititre™ FINMER plates with meropenem (Thermo Fischer, Diagnostics custom plates; YFÍNMER). APC148 was tested at fixed concentrations of 4, 8 and 16 µg/ml, alone, or in combination with avibactam at a fixed concentration of 8 µg/ml for strains carrying SBLs. The plates were incubated for 20 h at 37°C and results were interpreted as susceptible (S), susceptible, increased exposure (I) or resistant (R), in accordance with the EU-CAST clinical breakpoint table (20).

**Table 1:** Bacterial isolates included in this study, with sequence types (STs) from multilocus sequence typing (MLST) and molecularly characterized β-lactamases carried by each isolate, included.

| Species | Isolate/strain | Sequence type | β-lactamase(s) | Ref. |
| --- | --- | --- | --- | --- |
| *E. coli* | KresCPE0372 | ST2083 | NDM-5, OXA-181, CMY-42, TEM-1 | (21) |
| *K. pneumoniae* | KresCPE0353 | ST147 | NDM-1, KPC-2, CTX-M-15, OXA-1, SHV-11 | (21) |
| *E. coli* | #53 | ST95 | IMP-26, CTX-M-15, TEM-1 | (22) |
| *E. coli* | #36 | ST101 | NDM-7, CTX-M-15, OXA-1 | (22) |
| *E. coli* | #50 | ST410 | VIM-4, CTX-M-15, CMY-4, TEM-169 | (22) |
| *K. pneumoniae* | #22 | ST525 | NDM-1, OXA-181, SHV-11, CTX-M-15, OXA-1, TEM-1 | (22) |
| *K. pneumoniae* | K66-45 | ST11 | NDM-1, SHV-11, CTX-M-15, OXA1, OXA-9, TEM-1 | (22) |
| *K. pneumoniae* | K64-62 | ST2134 | VIM-1, SHV-12, TEM-1 | (22) |
| *K. pneumoniae* | BAA-1705<sup>TM</sup> | ART2008133 | KPC-2, SHV-1, TEM, OXA-18 | ATCC |
| *P. aeruginosa* | K34-7 | ST233 | VIM-2 | (23) |
| *P.aeruginosa* | PAN-64 | ST111 | VIM-2 | (24) |

**Table 2.** Zinc concentration in four different commercial brands of cation-adjusted Mueller-Hinton broth (caMHB), as determined by ITC-LC-MS (this study). The values were assessed with a 95% confidence interval. A nearly five-fold variation of the zinc concentrations was detected between the different caMHBs.

| caMHB manufacturer | Lot number | Total zinc concentration (µg/ml) | Total zinc concentration (µM) |
| --- | --- | --- | --- |
| Thermo Fischer (TF), T3462 | 584619 | 1.9 | 29.4 |
| Becton Dickinson (BD), 212322 | 313124 | 1.1 | 17.0 |
| Oxoid (OX), CM0405B | 3708386 | 0.6 | 9.2 |
| Millipore (MP), 90922 | 102445678 | 0.4 | 5.6 |

## Results

### Highly variable zinc concentration in different cation-adjusted Mueller Hinton broths

The zinc content in the four commercially available cation adjusted Mueller Hinton broths (caMHB) used for AST testing was determined, revealing considerable variation between brands (Table 2). The caMHB with the highest zinc concentration (Thermo Fischer) contained almost five-fold higher zinc compared to that of the caMHB with the lowest concentration (Millipore).

### Reduction in meropenem MICs by APC148 in different cation-adjusted Mueller Hinton broth brands with variable zinc concentrations

MIC experiments were then performed for meropenem alone and in combination with APC148, with a selection of *K. pneumoniae, E. coli* and *P. aeruginosa* isolates producing clinically relevant MBLs. The experiments were performed in caMHB from the four different manufacturers included in the study, and which had been shown to exhibit highly variable zinc content (Table 2). The MIC of meropenem alone was in the resistant (R) breakpoint range (MIC > 8 ug/mL) for all isolates included in the study. At 16 ug/ml, APC148 reduced the meropenem MIC to a susceptible (S) breakpoint category (≤ 2 ug/ml) for all isolates (Fig. 1). However, at lower concentrations of APC148 (8 or 4 ug/ml), the MIC of meropenem for the same isolates were highly dependent on the caMHB brand used in the assay, and was positively correlated (lower Zn, lower MIC) with the zinc content in the broth from the different manufacturers (Fig. 1). Thus, for the majority of isolates, when employing the two caMHBs with the lowest zinc concentrations (OX and MP; Table 2), APC148 at 4 or 8 ug/ml reduced the MIC of meropenem to susceptible levels. Corresponding MIC assays performed with the two caMHBs with the highest zinc concentrations (BD and TF) however, in the majority of cases led to meropenem MICs in combination with APC148 (at 4 or 8 ug/ml) of 32-64 ug/ml (Fig. 1), which would be interpreted as meropenem resistant by *in vitro* clinical diagnostics using the EU-CAST guidelines (20). As expected, for *K. pneumoniae* BAA 1705 which produces KPC-2 only, a serine-β-lactamase which exhibits a zinc-independent catalytic mechanism, MICs were not affected by either zinc content in the Mueller Hinton medium nor by the addition of APC148 (Fig. 1).

**Figure 1.**
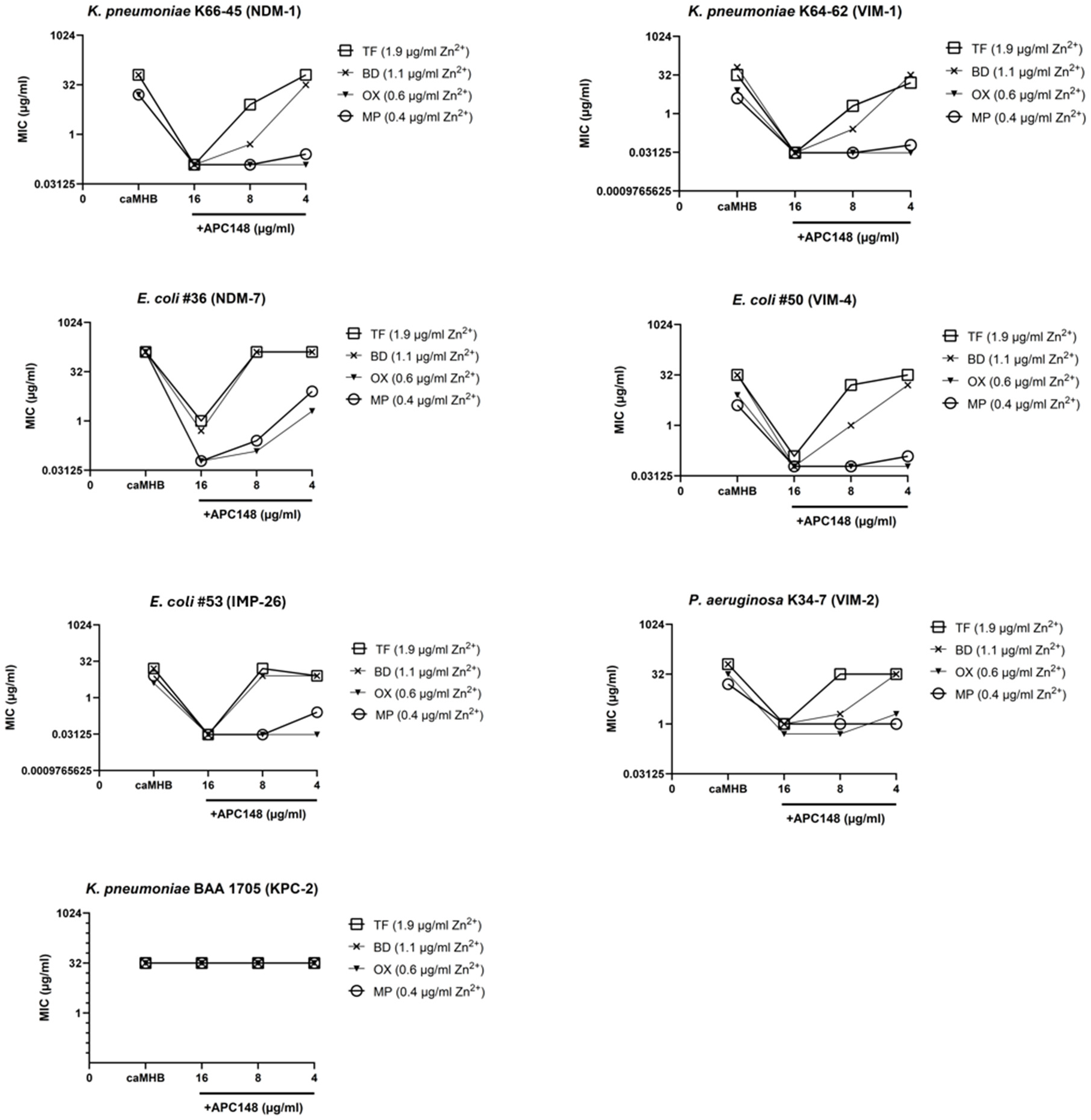
The MIC for meropenem alone (caMHB), or in combination with APC148 at a fixed concentration of 4, 8 and 16 ug/ml, respectively, against clinical isolates of *E. coli, K. pneumoniae*, and *P. aeruginosa*, in four different brands of caMHB with varying zinc content (0.4 – 1.9 μg/mL; TF: Thermo Fisher, MP: Millipore, BD: Beckton Dickinson, OX: Oxoid). *K. pneumoniae* BAA 1705 (does not carry MBL) was included as a negative control. Each datapoint represents the modal MIC of three replicate experiments.

### Reduction in meropenem MICs by APC301 in different cation-adjusted Muller Hinton broth brands with variable zinc concentrations

The effect on APC148 performance of employing different MHB brands with variable zinc concentrations was similarly tested on clinical isolates of Gram-negative bacteria (*K. pneumoniae, E. coli, P. aeruginosa*) carrying different variants of both MBL- and SBL-encoding genes. Here, APC148 was used in combination with meropenem and the serine-β-lactamase (SBL) inhibitor avibactam (APC301, triple combination). The MICs of meropenem alone were in the resistant (R) breakpoint range (MIC > 8 ug/mL) for all isolates tested. Addition of avibactam alone had no notable effect, except for with *K. pneumoniae* KresCPE0353 (NDM-1, KPC-2), where a small MIC reduction could be observed in the caMHBs with the lowest zinc content (Fig. 2).

**Figure 2.**
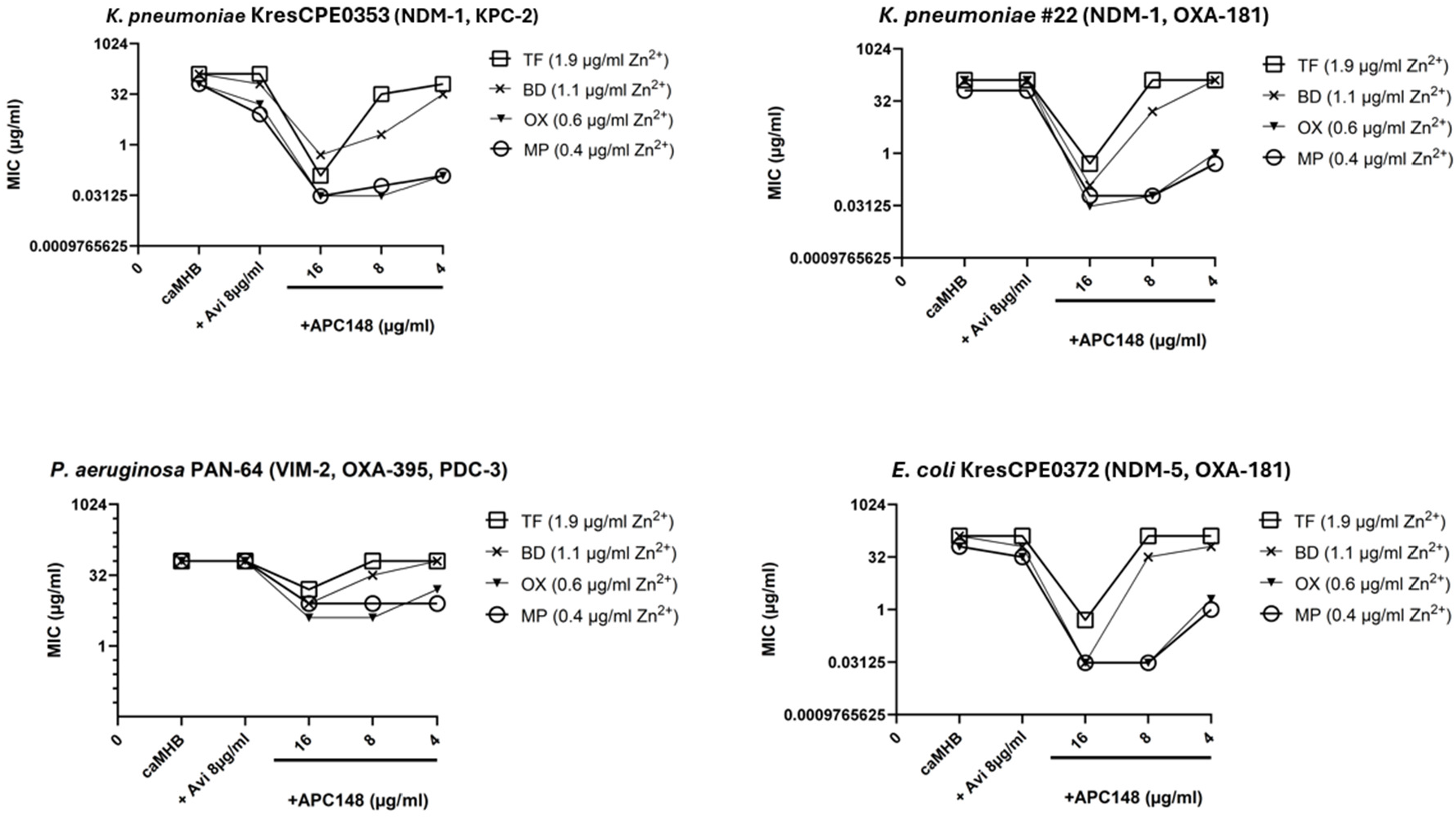
Meropenem MICs with meropenem alone (caMHB) or in combination with avibactam (8 ug/ml; +Avi), and avibactam (8 ug/ml) and APC148 (+APC148) at three different concentrations (16ug/ml, 8 ug/ml and 4 ug/ml, respectively), for four strains of *E. coli, K. pneumoniae*, and *P. aeruginosa* producing both MBL and SBL enzyme(s) in different combinations. Experiments were performed in caMHB from four different manufacturers, spanning zinc concentrations from 0.4 ug/ml to 1.9 ug/ml (as indicated; TF: Thermo Fisher, MP: Millipore, BD: Beckton Dickinson, OX: Oxoid). Each data point represents the modal MIC of three replicate experiments.

By addition of APC148 at 16 µg/ml in combination with avibactam (8 µg/ml), the meropenem MIC was brought down to susceptible (S) levels (0.5 – 0.03) for all *K. pneumoniae* and *E. coli* stains, regardless of zinc content of the caMHB (Fig. 2). Reducing the APC148 concentration to 4 or 8 ug/ml (in combination with 8 µg/ml avibactam), MICs in the susceptible range (S) were retained only when using the two caMHBs with the lowest zinc concentrations. Employing the two caMHBs with the highest zinc concentrations on the contrary led to meropenem MICs of 32-64 ug/ml (Fig. 2). For *P. aeruginosa* PAN-64, a similar pattern was observed, where at 16 µg/ml APC148 and 8 µg/ml avibactam, the meropenem MICs were reduced for all caMHBs, although only to the susceptible, increased exposure (I) category (4-8 ug/ml). At 4 and 8 µg/ml APC148, this category was retained only when employing the two MHBs with the lowest zinc content (Fig. 2).

## Discussion and conclusions

In this study we have focused on the effect of the varying zinc content of commercially available Mueller Hinton broths and its effect on the outcome of *in vitro* AST testing with the new chelator-based metallo-*β*-lactamase inhibitor APC148. At infection sites, zinc concentrations have been shown to be in the nanomolar range (15). It has thus been argued that low zinc conditions during *in vitro* testing may potentially better reflect the *in vivo* situation during infection (6). APC148 has previously been tested *in vivo* in various models, including a murine neutropenic peritonitis model. Subcutaneous treatment with meropenem (33 mg/kg) and APC148 (10 mg/kg) in combination resulted in a significantly lower live cell count (CFU/mL) of a meropenem-resistant NDM-1-producing *K. pneumoniae* strain in both peritoneal fluid (*P* < 0.0001) and in blood (*P* < 0.01), as compared to treatment with meropenem alone (18).

For the *in vitro* experiments perfomed here, while at 16 µg/ml APC148 lowered the MIC in all caMHBs, however the zinc content of the growth medium had a marked effect on meropenem MIC reduction at concentrations of APC148 lower than 16 µg/ml. This may be attributable to zinc in the caMHB saturating the environment and thus attenuating the zinc-chelating effect of APC148. Previous studies have shown the importance of zinc content when determining the susceptibility of MBL-producing isolates in routine diagnostics (5-7, 15, 17). The results from the present study are in accordance with recent publications (16, 17) stating that there is considerable variability in zinc content among commercial caMHBs, potentially resulting in variability in SIR classifications of isolates across laboratories due to use of MHB from different manufacturers. Clearly, as we show in the current study, this variation in zinc content can affect SIR interpretations during routine diagnostics AST testing with MBL inhibitors like APC148, which employ zinc chelation as its mechanism of action. Thus, the results presented in the current study clearly demonstrates the need to employ APC148 at a minimum concentration of 16 µg/ml during *in vitro* AST testing by broth microdilution, in order to avoid issues of brand-dependent effects from the caMHB used. This applies both for bacterial isolates producing MBL as the main resistance mechanism, as well as for isolates that are dual MBL and SBL producers, when using APC148 in combination with the SBL inhibitor avibactam.

## Acknowledgements

This research was funded by a grant from The Research Council of Norway through the IPN programme (project number 346300, RESTORE). We gratefully acknowledge Ørjan Samuelsen at the Norwegian National Advisory Unit on Detection of Antimicrobial Resistance for providing clinical bacterial isolates used in this study.

